# Development of heat shock resistance in *L. pneumophila* modeled by experimental evolution

**DOI:** 10.1101/2023.04.27.538606

**Authors:** Jeffrey Liang, Sebastien P. Faucher

**Affiliations:** Department of Natural Resource Sciences, McGill University, Sainte-Anne-de-Bellevue, Québec, Canada; Centre de Recherche en Infectiologie Porcine et Avicole (CRIPA), Faculté de Médecine Vétérinaire, Université de Montréal, Saint-Hyacinthe, QC, Canada

**Keywords:** Legionella pneumophila, Hot water distribution systems, Adaptive laboratory evolution, Experimental evolution, Heat resistance, Pasteurization

## Abstract

Because it can grow in buildings with complex hot water distribution systems (HWDS), healthcare facilities recognize the waterborne bacterium *Legionella pneumophila* as a major nosocomial infection threat and often try to clear the systems with a pasteurization process known as superheat-and-flush. After this treatment, many facilities find that the contaminating populations slowly recover, suggesting the possibility of *in situ* evolution favouring increased survival in high temperature conditions. To mimic this process in a controlled environment, an adaptive laboratory evolution (ALE) model was used to select a wild-type strain of *L. pneumophila* for survival to transient exposures to temperatures characteristic of routine hot water use or failed pasteurization processes in HWDS. Over their evolution, these populations became insensitive to exposure to 55 °C and innovated the ability to survive short exposures to 59 °C heat shock. Heat-adapted lineages maintained a higher expression of heat shock genes during low-temperature incubation in freshwater, suggesting a pre-adaptation to heat stress. Although there were distinct mutation profiles in each of the heat-adapted lineages, each acquired multiple mutations in the DnaJ/DnaK/ClpB disaggregase complex, as well as mutations in chaperone *htpG* and protease *clpX.* These mutations were specific to heat shock survival and were not seen in control lineages included in the ALE without exposure to heat shock. This study supports *in situ* observations of adaptation to heat stress and demonstrate the potential of *L. pneumophila* to develop resistance to control measures.

**Importance:** As a bacterium that thrives in warm water ecosystems, *Legionella pneumophila* is a key factor motivating regulations on hot water systems. Two major measures intended to control *Legionella* are the maintenance of high circulating temperatures to curtail growth and the use of superheat-and-flush pasteurization processes to eliminate established populations. Although hospitals are particularly vulnerable to nosocomial pneumoniae caused by *Legionella*, they recurrently experience recolonization of their hot water systems after treatment. To understand these long-term survivors, we have used an experimental evolution model to replicate this process. We find major differences between the mutational profiles of heat-adapted and heat-naïve *L. pneumophila* populations, including mutations in major heat shock genes like chaperones and proteases. This model demonstrates the value of appropriate heat treatment of *L. pneumophila* contaminated systems and – in an analogue to antibiotic resistance – the importance of complete eradication of the resident population to prevent selection for more persistent bacteria.

## Introduction

As an obligate intracellular pathogen, the bacterium *Legionella pneumophila* relies on infection of eukaryotic cells in order to acquire nutrients and replicate (1). Though it usually lives in freshwater ecosystems where it targets free-living eukaryotic hosts from diverse clades, *L. pneumophila* is now also known to infect humans where it targets alveolar macrophages (2). Its contamination of human-built infrastructure favours infection by aerosolizing the bacteria and facilitating its travel into the lungs (2, 3). Globally, *L. pneumophila* is the main cause of Legionnaires’ Disease, a life-threatening pneumonia whose severity and under-diagnosis have led to its high overall case-fatality rate, estimated at between 5 and 10%, with even higher death rates for nosocomial cases (4–6). Early suspicions focused on dispersal through contaminated cooling tower emissions (7, 8), but *L. pneumophila* has since been found in diverse engineered water systems, including in whirlpool spas, decorative fountains, and – significantly for nosocomial infection – in hospital plumbing systems (9).

The engineering of water systems has a major impact on the growth potential of *L. pneumophila* (10, 11). Large buildings with complex plumbing systems are particularly vulnerable to contamination, and studies have shown that multiple clades of *L. pneumophila* can stably co-exist in these systems (12). As a mesophilic bacterium which replicates best between 20 and 45 °C (13), *L. pneumophila* favours colonization of hot water distribution systems (HWDS). Deprecated fixtures, as well as the improper design of plumbing networks or incorrect estimation of water demand, can lead to stagnation and the existence of dead legs. These regions of low water flow allow scale and biofilms to accumulate – physical structures that can physically protect *Legionella* from exposure to disinfectants (14, 15). Policies that decrease water temperatures in favour of energy savings can result in tepid temperatures that favour the growth *L. pneumophila* and of its eukaryotic hosts (16, 17). Hospitals are particularly at risk both because they must balance hot water temperatures against the risks of scalding and because they often house immunocompromised patients who are particularly threatened by exposure to recurrent bacterial contaminants (18, 19).

Along with chemical treatment via chlorination or copper-silver ionization, the maintenance of high-temperatures along the length of a plumbing system from the water heater through to the terminal outlets is a critical control measure against *L. pneumophila* (20). Remedial high-temperature pasteurization processes, collectively termed superheat-and-flush, aim to push hot water temperatures above what *L. pneumophila* can tolerate (21, 22). The implementation of these strategies can differ depending on the specific architecture of the HWDS, but they aim to raise temperatures throughout the entire system through to the distal outlets. Though remedial pasteurization can be effective at the short-term clearance of *Legionella*, longitudinal studies have shown that re-contamination is common over month or year-long time scales (23–25). It has been reported that repeated use of high-temperature pasteurization to disinfect these water systems can instead lead to the local evolution of more heat resistant *L. pneumophila* populations, like the familiar phenomenon of antibiotic tolerance (20, 22, 23).

In bacteria, heat stress can damage several vital subcomponents of the cell by causing membrane depolarization and damage, destabilization of the nucleoid, and the inactivation and aggregation of vital enzymes and structural proteins (26–28). Bacteria have developed a sophisticated heat shock response, centralized through the alternative sigma factor RpoH that redirects transcription towards stress resistance by driving the production of heat shock proteins (29, 30). These heat shock proteins buffer the toxic effects of misfolded and aggregated proteins in the cell – either by promoting refolding or by targeting them for destruction (31, 32). These proteins are broadly conserved among bacterial species and include proteases as well as chaperones, such as HtpG and DnaK, and their co-chaperone proteins, such as DnaJ and GrpE. After the thermal shock is resolved, these proteins are also involved in a negative feedback loop targeting RpoH for destruction by FtsH, a membrane-bound protease (33, 34). This organization is conserved in *L. pneumophila*, with five heat shock proteins identified in radiolabelling studies (35) and many more annotated from the published genetic sequence (36, 37). Heat shock proteins are also involved in the early pathogenesis in *L. pneumophila* by promoting the internalization of the bacterium by the host cell and reorganizing host trafficking processes (38–40). The LetAS and CpxRA two-component systems are also known to influence heat shock survival, likely mediated through their regulation of life phase switching during growth and starvation (41, 42).

Adaptive laboratory evolution (ALE) is an experimental concept that manipulates a chosen organism in a laboratory setting to providing insights into complicated phenomena from the wild (43, 44). To simulate long-term exposure to high temperatures in HWDS, we can depend on the short generation times and the relative ease of culture and storage of *L. pneumophila* in laboratory conditions to study the resulting evolution in a controlled and repeatable process. ALE has been applied in numerous bacterial and non-bacterial species to study the trajectories and repeatability of evolutionary responses without *a priori* expectations on the loci which will be targeted (45, 46). Prior studies have used ALE to adapt *Escherichia coli* to growth in stably high-temperature or low pH conditions, as well as to survival acute population bottlenecking by ionizing radiation (47–49). Notable ALE experiments conducted with *L. pneumophila* have been used to study resistance to fluoroquinolone and macrolide antibiotics (50, 51) and adaptation to mouse macrophages (52). As heat stress non-specifically challenges multiple components of the bacterial cell, ALE makes an appropriate protocol to study the range of mutations driving the increasing heat tolerance of *L. pneumophila* populations observed in hot water systems.

To bridge the gap between *in situ* observations of *L. pneumophila*’s adaptation to pasteurization in HWDS and an understanding of its underlying genetic mechanisms, we tailored an ALE study design to investigate the evolutionary paths that the environmental pathogen can take to develop a tolerance of heat shock. The aim was to document the genetic changes that surviving population would develop under transient exposure to high temperatures, simulating failed pasteurizations or bursts of hot water from routine outlet use. We report that ALE can reliably induce resistance to heat shock in adapted lineages through underlying mutations that cause differences both in transcriptional regulation and heat shock protein activity. In our model, adaptation to heat shock resulted in no fitness deficits during axenic or infectious growth, supporting epidemiological studies that connect HWDS isolates to clinical isolates from exposed individuals.

## Results

Studies tracking the population structure over time of *L. pneumophila* growing in HWDS have repeatedly confirmed that persistence and microevolution is more common than recontamination from the incoming water supply. Even more troubling, repeated exposure to high temperatures through bursts of hot water demand or failed cycles of superheat-and-flush are liable to favour heat resistance in the surviving bacteria. To simulate this process and study it in a controlled environment, we designed an adaptive evolution system to simulate transient and sublethal exposures to high temperatures. Based on pilot data and typical global public health recommendations (13), we exposed the study populations to 55 °C by immersion in a circulating water bath for 15 minutes, as described in Materials and Methods. We maintained six replicate lineages of the wild-type Philadelphia-1 strain to determine the replicability of our evolutionary pathways under heat shock (Figure 1A). In parallel, we also maintained six replicate control lineages to compensate for mutations that favour growth in laboratory conditions and isolate the changes related to heat shock (Figure 1B). Bacterial population sizes were bottlenecked at each cycle by either selection for survivors of heat shock or by sub-sampling of control populations suspended in freshwater.

**Figure 1.**
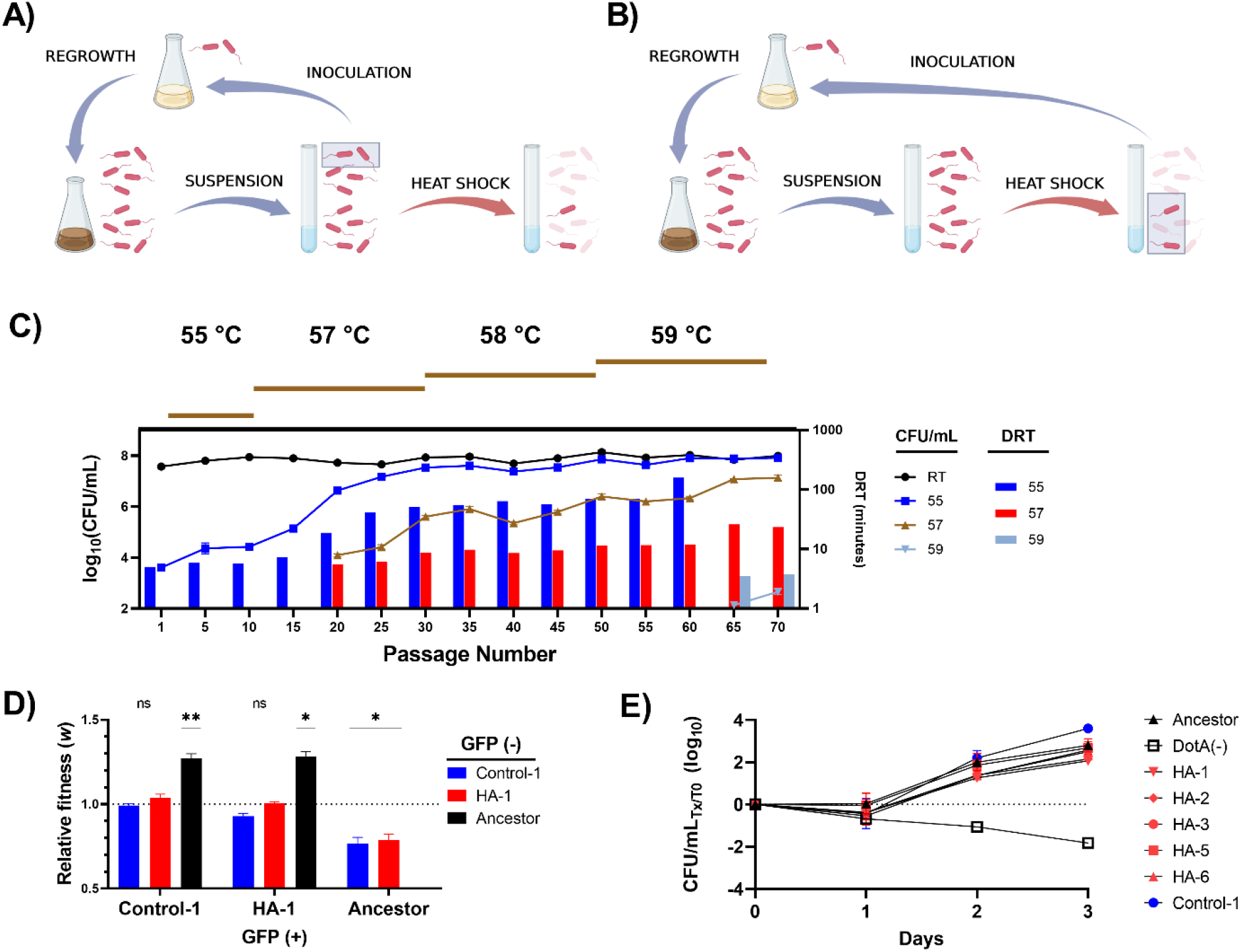
ALE increased the resistance of *L. pneumophila* to heat shock. The organization of the ALE protocol, contrasting the heat-adaptation (A) and control (B) branches (Created with BioRender). Each panel depicts one of 70 cycles, with highlighted rectangles showing the population bottleneck for each branch. In brief, after collection from post-exponential growth in AYE and 24-hour suspension in Fraquil, control lineages were propagated by subsampling the suspended population to re-inoculate AYE while heat-adapted lineages were propagated by re-inoculating AYE with the population surviving heat shock. C) Survival of heat-adapted *L. pneumophila* lineages following 20 minutes of exposure to 55, 57, and 59 °C heat shock for 20 minutes (lines, left axis). Decimal reduction times (DRT) were calculated for each temperature and are plotted for each tested passage (right axis, bar; omitted for passages 65 and 70 at 55 °C because there was an insignificant level of cell death). D) Direct competition between green-fluorescent isolates and non-fluorescent isolates growing in AYE. Significance calculations show one-sample *t*-test against 1.0 (neutral relative fitness), *n* = 3. E) Infection and replication within *V. vermiformis* in modified PYNFH without FBS, *n* = 3. The avirulent DotA(-) strain is used as a negative control.

Over the 70 cycles of the ALE protocol, we raised the heat shock temperature on an *ad hoc* basis to maintain a stable level of population loss in the heat adapted lineages from 55 °C at the beginning of the experiment to 59 °C at its conclusion (Figure 1C). The resulting six heat-adapted populations were named HA-1 through HA-6 and the six control lineages were named C-1 through C-6 (Table 1). Whole genome sequencing revealed that HA-4 had been contaminated by HA-6 by the end of the experiment, so it was removed from final analysis. Regardless, all replicate heat adapted lineages showed a robust increase in heat shock resistance when challenged by heat shock for 20 minutes (Figure 1C). The decimal reduction time required to reduce the initial populations by 90% increased from 303 seconds at 55 °C at the outset of the ALE to a minimum of 9546 seconds by passage 6 (Figure 1C). After 65 and 70 cycles of selection, there was no significant population drop even after a 30-minute exposure. After 20 and 65 passages respectively, populations could survive 57 °C and 59 °C for 20 minutes. In contrast, neither the control lineages nor the ancestral strain was able to tolerate 57 °C for 20 minutes (data not shown).

**Table 1:** Bacterial strains and plasmids used in this study. Except where noted, all strains used were of *L. pneumophila*

| Strain | Description | Source |
| --- | --- | --- |
| Philadelphia-1 | Philadelphia 1976 | ATCC33152 |
| KS79 | JR32 ΔcomR | (107) |
| ΔhtpG | KS79 htpG::Kn<sup>pSF6</sup> | (75) |
| htpG<sup>G83E</sup> | KS79 htpG::htpG<sup>HA-3</sup> + Kn pGEMT-easy | This study |
| htpG<sup>Q148*</sup> | KS79 htpG::htpG<sup>HA-5</sup> + Kn pGEMT-easy | This study |
| htpG<sup>Δ1bp nt603</sup> | KS79 htpG::htpG<sup>HA-6</sup> + Kn pGEMT-easy | This study |
| *E. coli* DH5α | supE44 ΔlacU169 (Φ80lacZΔM15) hsdR17 recA1 endA1<br>gyrA96 thi-1 relA1 | Invitrogen |
| Plasmid | Description | Source |
| pMMB207c | RSF1010 derivative, IncQ, lacIq, Cm<sup>r</sup>, Ptac, oriT, ΔmobA | (108) |
| pXDC39 | pMMB207c ΔPtac, ΔlacI, Cmr | Xavier Charpentier |
| pSF6 | pGEMT-easy-rrnB | (109) |
| pXDC31 | pMMB207C + *gfp* | (110) |
| phtpG<sup>G83E</sup> | pXDC39 + htpG<sup>HA-3</sup> | This study |
| phtpG<sup>Q148*</sup> | pXDC39 + htpG<sup>HA-5</sup> | This study |
| phtpG<sup>Δ1bp nt603</sup> | pXDC39 + htpG<sup>HA-6</sup> | This study |
| phtpG<sup>WT</sup> | pXDC39 + htpG<sup>Philadelphia-1</sup> | (75) |

Despite their acquired ability to tolerate heat shock, the heat adapted lineages did not lose the ability to grow axenically or in host cells. We directly competed the evolved lineages against their ancestral strain by growing them together in co-culture to test for relative fitness. Plasmids pXDC31 and pMMB207c, which differ only in that pXDC31 has an IPTG-inducible *gfp* insert, were used to distinguish between strains. Both plasmids were electroporated into the Philadelphia-1 ancestor as well as Passage 70 populations of both HA-1 and C-1 and transformed isolates were used in the competition experiments. To control for possible fitness confounds introduced by the plasmid inserts, all pairs were tested with a reciprocal expression of *gfp* in both competitors (e.g., HA-1 pXDC31 was competed against Philadelphia-1 pMMB207C and HA-1 pMMB207C was competed against Philadelphia-1 pXDC31). Control competitions of HA-1 and C-1 with themselves showed a neutral relative fitness, as expected (*p* = 0.4123, 0.5233). Both ALE-derived competitors were also at neutral relative fitness against each other (*p* = 0.0547, 0.2113), indicating that both were equally well adapted to growth in AYE. In contrast, Philadelphia-1 was significantly less fit than either HA-1 (*p* = 0.0120, 0.0235) or C-1 (*p* = 0.0090, 0.0244), supporting our hypothesis that the populations would evolve better growth in AYE in tandem with heat resistance. To test whether this increased growth potential in AYE was replicated during host cell infection, we tested for three-day bacterial yield after growth in the amoeba *V. vermiformis*. Consistent with a specific adaptation to growth in cell-free conditions, replication of the heat-adapted lineages did not significantly differ from those of Philadelphia-1 or the control lineage (Figure 1E).

Because the early bacterial heat shock response relies largely on a re-direction of transcription by the *rpoH* sigma factor, we expected that there would be a difference in RNA expression profiles between our heat-adapted, control, and ancestral strains. To represent the regulators and effectors of the transcriptional heat shock response, we measured differential expression of three sigma factors and 7 heat shock proteins. These were chosen to represent genes likely to be involved in the heat shock response, as well as those mutated in multiple lineages under heat shock selection. Upon heat shock, the ancestor Philadelphia-1 displays significant up-regulation in *clpB*, *dnaJ*, *dnaK*, *groL*, and *htpG* expression but not of *clpX*, *lon*, or *phaP* by two-way ANOVA (Figure 2 A). Similarly, *rpoH* is the only sigma factor whose RNA levels are elevated (Figure 2B). Compared to Philadelphia-1, the heat-adapted lineages (and to a lesser extent, the control lineages) over-express heat shock protein mRNAs prior to heat shock exposure (Figure 2C). This over-expression is no longer seen when comparing expression profiles after heat shock, suggesting a pre-adaptation to thermal stress in chronically heat-exposed populations (Figure 2D).

**Figure 2:**
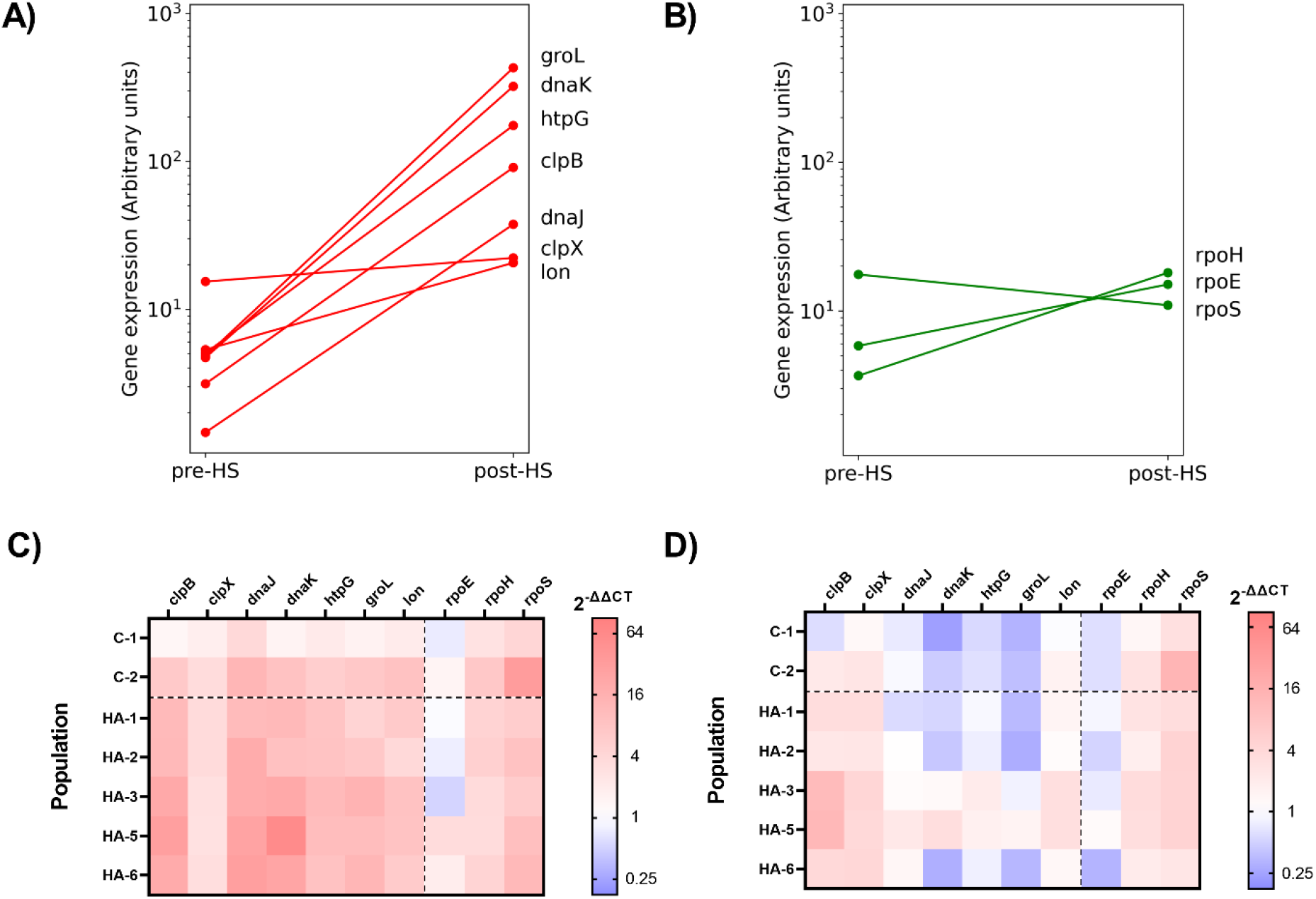
qPCR analysis of RNA expression levels in the ancestor Philadelphia-1 following 5 minutes of exposure to 55 °C heat shock for heat-shock response genes (A) and sigma factors (B). Ct values were normalized using 16S rRNA levels and converted to arbitrary expression levels, *n* = 3. Relative difference in expression in RNA samples collected before heat shock (C) and after heat shock (D) between sampled populations (C, control lineages; HA, heat-adapted lineages) and Philadelphia-1, *n* = 3.

To identify the mutations acquired after 70 cycles of the ALE, we used Breseq, a pipeline designed for the analysis of short read data from bacterial evolution experiments. To capture the population diversity, we isolated 10 clones from each heat-adapted lineage and 5 clones from each control lineage for whole-genome sequencing. As described in Materials and Methods, per-isolate analysis of the sequence reads showed gaps of missing coverage, so reads from the replicate clones of each lineage were pooled and re-analysed in the polymorphism setting of Breseq to measure allele frequency of novel mutations. Re-sequencing of our Philadelphia-1 strain showed that it differs from the published genome (CP015927.1) by the absence of a 38 kbp island pPh38 and by a missense mutation in the *letA* gene (Table 3). Genes mutated in replicate heat-adapted lineages included multiple heat shock genes (*htpG*, *clpB*, *clpX*, *dnaJ*, and *dnaK*), as well as lpg0563 (*phaP*), lpg2351 (*mavF*), and genes associated with cell wall synthesis (*mreC* and *rodA*) (Figure 3 and Table 3). A separate mutation profile was seen in the replicate control lineages, with multiple hits in *nusG*, *rpoB*, *cpxR*, and *lidA* (Figure 3). Each lineage displayed a unique mutational profile, showing that *L. pneumophila* can follow multiple evolutionary pathways to acquire strong heat resistance (Supplementary table 1).

**Figure 3:**
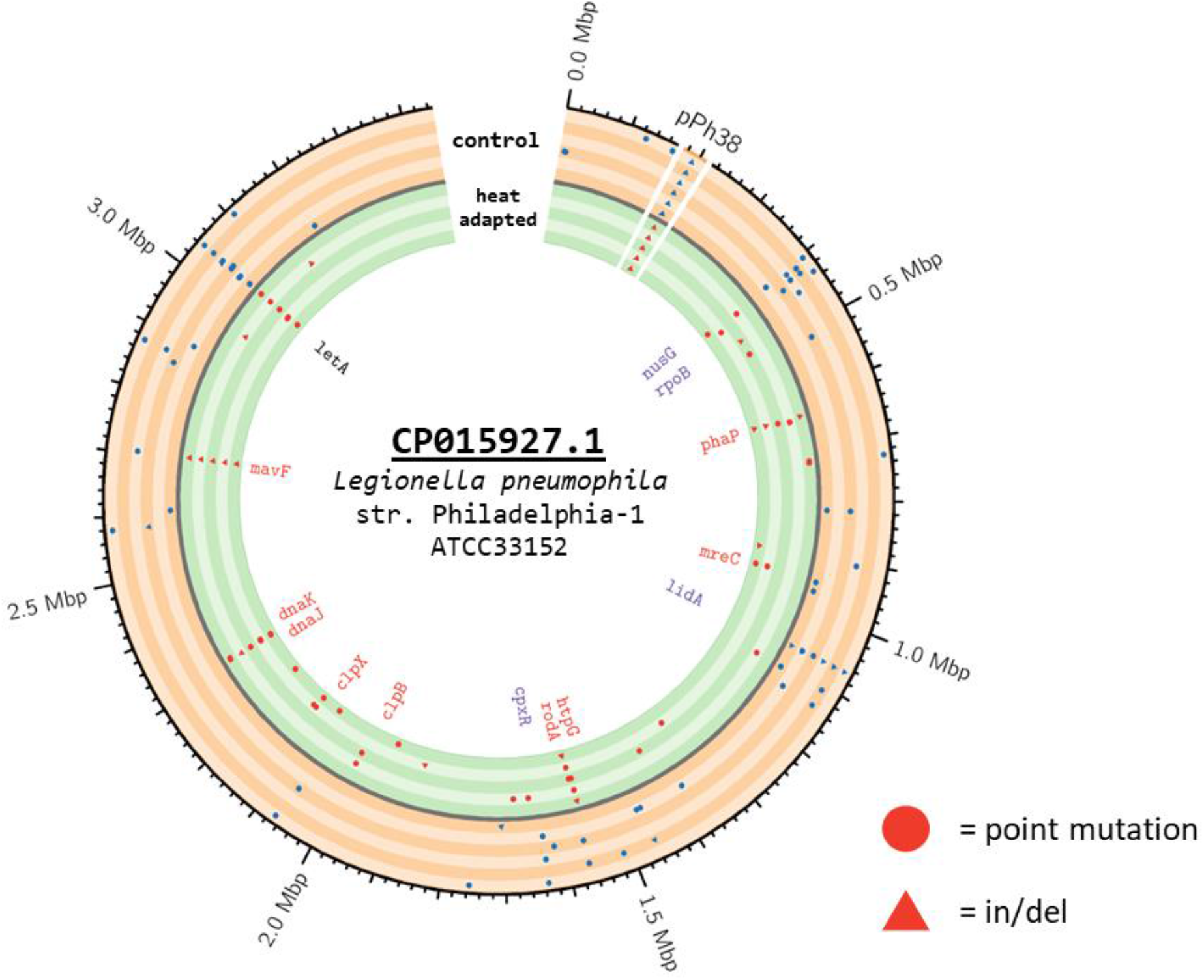
Derived mutations identified with Breseq at a frequency of at least 25% in replicate control and heat-adapted lineages aligned to the genome of the ancestral strain Philadelphia-1 (106). The outermost concentric circle shows the single chromosome of *L. pneumophila*, as well as a reversibly-integrated pPh38 element which was not retained in our lab strain. Each band shows the mutations observed across control lineage (orange) and heat-adapted (green), with circles representing point mutations and triangles representing insertions and deletions. Genes of interest in the interior are labelled by colour with red genes mutated in multiple heat-adapted lineages and blue genes mutated in multiple control lineages.

The *L. pneumophila* homologue of *htpG* (bacterial 90 kilodalton heat shock protein) acquired mutations in three of the heat-adapted lineages including a nonsense mutation in HA-5 (Q148*) and a frameshift mutation in HA-6 (del1bp 503). Sequencing of HA-4 after 30 passages (before it had been contaminated) also showed a third distinct loss-of-function mutation (del1bp 694) (Supplementary table 2). Though surprising for a known heat shock protein, this was consistent with our prior finding that a deletion mutant lacking *htpG* showed increased survival under heat stress. Western blotting confirmed that the typical HtpG protein was not produced in HA-5 or HA-6, while the missense mutation (G83E) in HA-3 did not affect antibody binding (Figure 4A). The *htpG* alleles found in HA-3, HA-5 and HA6 were mobilized into the chromosome of the lab strain KS79 by allelic exchange in tandem with a selectable marker. Phenotypically, the mutant alleles of *htpG* raised heat shock resistance in *L. pneumophila* to a similar magnitude as the previously constructed full deletion mutant (Figure 4B). Reciprocally, none of the mutant alleles were able to complement the raised heat resistance of the deletion mutant (Figure 4C).

**Figure 4:**
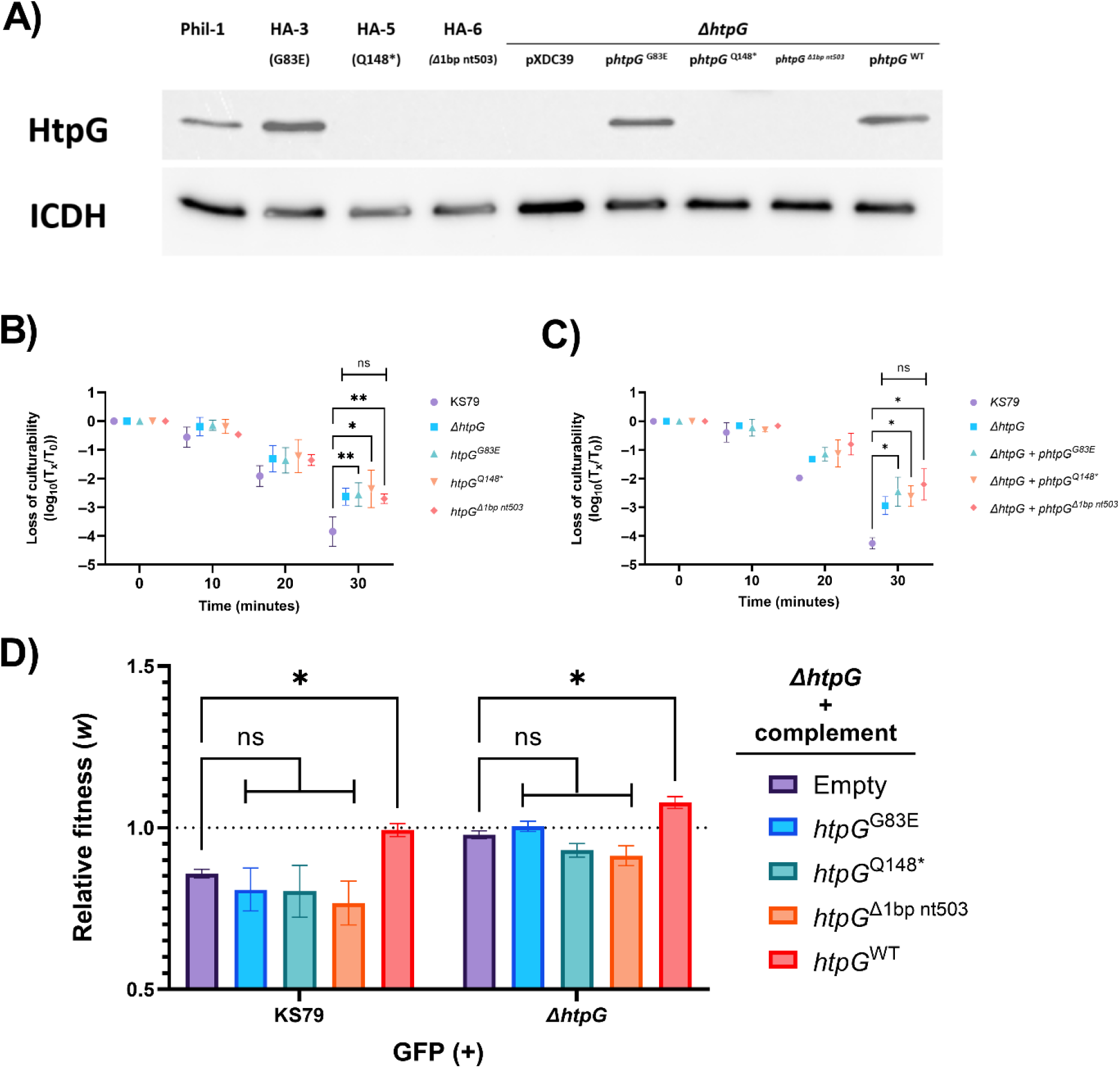
A) Anti-*htpG* Western blots visualizing the expression of HtpG in lineages HA-3, 5 and 6 and in constructed merodiploid mutants. B) Survival at 55 °C of knock-in exchange mutants expressing derived alleles of *htpG* in place of the wild-type allele, *n* = 3. C) Survival at 55 °C of an *htpG* deletion mutant heterologously expressing different alleles of *htpG* found in treated lineages, *n* = 3. D) Relative fitness of the *htpG* deletion mutant complemented with an empty plasmid, derived, or ancestral *htpG* against the wild-type and deletion mutant strains.

**Table 2:**
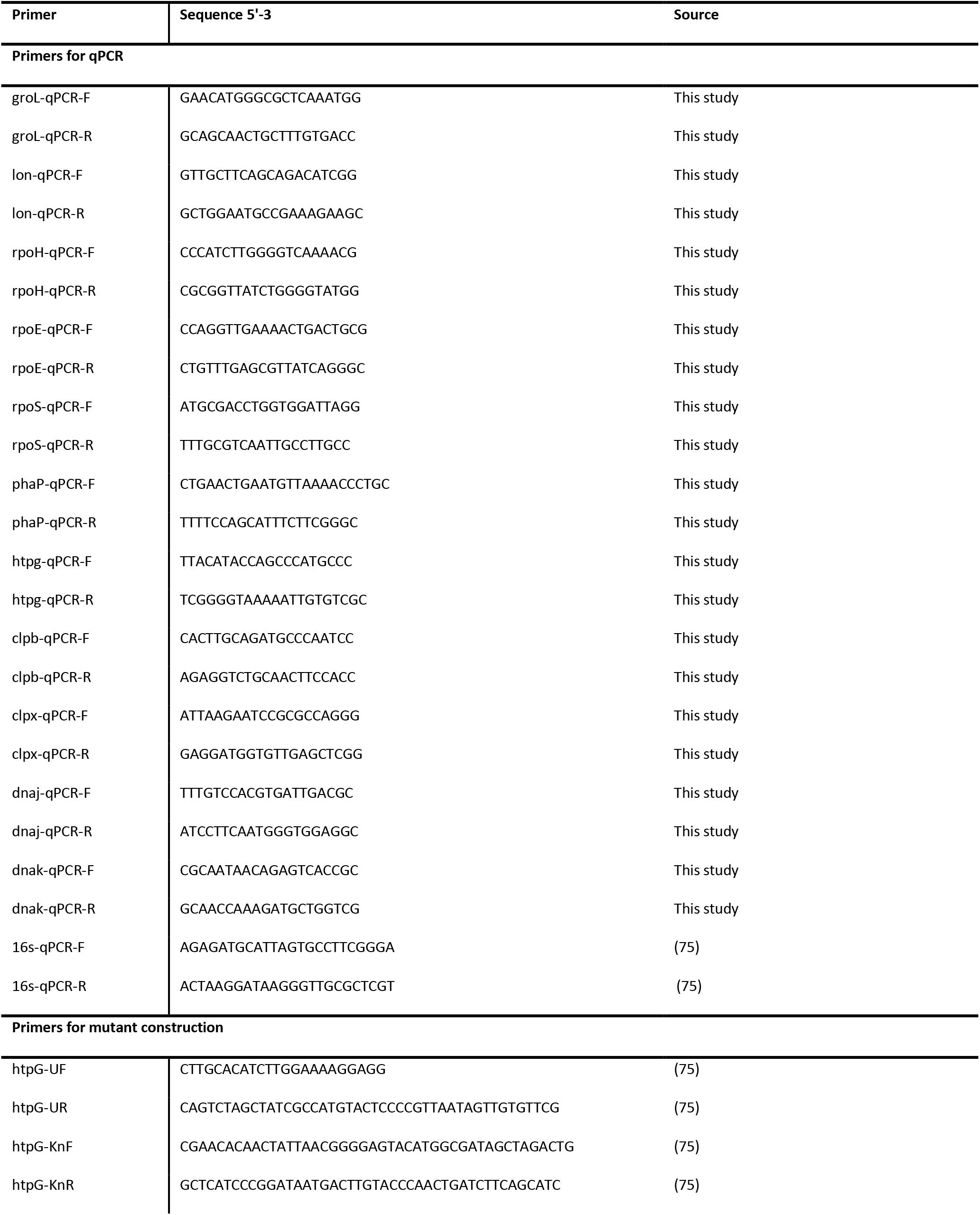
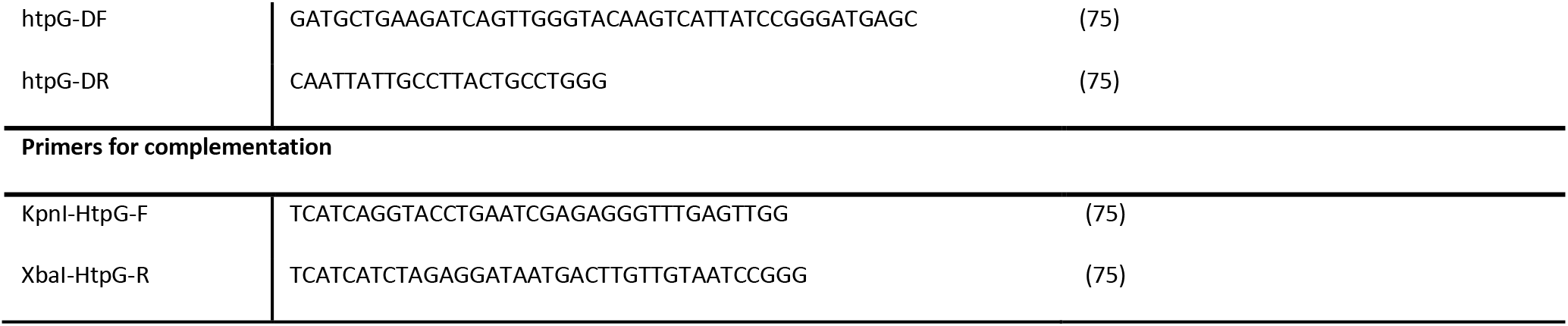
All primers used during this study

To test whether *htpG* mutation was favoured only due to its role in heat shock, we constructed a competition assay to compare the growth of KS79 and of the deletion mutant against the mutant harbouring the different alleles of *htpG* (Figure 4D). The wild-type KS79 under-competed against any strain containing a blank plasmid or plasmid carrying any derived allele of *htpG*. Conversely, it was neutrally fit against the *htpG* mutant complemented with a wild-type allele of *htpG*. KS79 and the deletion mutant both grew significantly better in the presence of competitors harbouring the ancestral allele of *htpG* compared to competitors harbouring a blank plasmid, suggesting that HtpG, in addition to conferring reduced heat shock resistance, may also reduce growth potential in AYE.

## Discussion

### *L. pneumophila* rapidly adapts to repeated exposure to heat shock

As an opportunistic colonizer of plumbing infrastructure, *L. pneumophila* is commonly recognized as a major hazard under the purview of hospital facilities managers (18). To model its long-term residence in hot water systems, our system of laboratory evolution was able to robustly raise the heat tolerance levels of wild-type *L. pneumophila*. The lineages’ survival of 55 °C – a commonly recommended circulation temperature (53, 54) – highlights the scale of their adaptation. Prior to their passage through the ALE model, the ancestral strain could survive only 5 minutes of exposure to this temperature before suffering a population reduction of 90% (Figure 1C). After 65 rounds of selection, the heat-adapted lineages could sustain exposure to the same temperature for at least 30 minutes without showing significant population loss. The final selection condition – 15 minutes at 59 °C – was much more intense that what the ancestral strain could survive and indeed our control lineages, which were also tested for heat shock survival, consistently failed to survive temperatures of 57 °C and higher.

A major concern underlying our experiment was that *L. pneumophila*, accustomed to replication inside eukaryotes, would simultaneously favour adaptation towards growth in liquid media and lose the ability to replicate inside host cells. We included replicate control lineages to anticipate the changes that might be seen after repeated growth in AYE. Testing populations from both branches of the evolution experiment, we found that both branches of the experiment retained their ancestral ability to replicate inside *V. vermamoeba* (Figure 1E). On the other hand, passage in axenic culture significantly elevated the relative fitnesses of both the control and heat-adapted lineages in AYE culture compared to Philadelphia-1 (Figure 1D). Despite relaxed constraints during the selection of the control lineage, the heat-adapted lineage had an equal ability to grow in AYE or inside host amoeba without needing to trade-off the ability to tolerate heat shock.

### Tandem adaptation of transcription and translation during adaptation to axenic culture

Consistent with their separate evolutionary pressures, the control and heat-adapted lineages adopted divergent evolutionary patterns. RNA transcription genes *nusG* and *rpoB* (55, 56) were both non-synonymously mutated in multiple control lineages, as was the S10 ribosomal protein *nusE* (Table 3 and Figure 3). RpoB is the beta-subunit of the RNA polymerase (RNAP) complex and is commonly mutated in bacterial evolution studies in diverse contexts including acid resistance, ionizing radiation, and atmospheric pollution (47, 49, 57). Four different missense mutations in *rpoB* were independently fixed in two control lineages (G759E and N1350T) and in two heat-adapted lineages (G138E and K924N). The transcription factor NusG binds to the RpoB through its N-terminal domain and to NusE through its C-terminal domain, coupling transcription to translation (55, 58). The two mutations seen in NusG (R136I and V143D) both sit within the C-terminal KOW domain (59) and are in lineages C-2 and C-6 – mutually exclusive with mutations in *rpoB* and *nusE* harboured in lineages C-4 and C-5. NusG enhances both RNAP processivity and rho-dependent termination, so the partition of these mutations could suggest incompatibility between the NusG and RpoB/NusE mutations, though epistasis was not tested in this study (55, 56, 60). From ALE experiments in *E. coli,* mutations in *rpoB* are known to cause broad changes to global transcription (61) and can optimize replication rates during exponential growth (62, 63). In these experiments, the same mutations in *rpoB* that contribute to faster exponential growth rates also reduce the ability of these strains to tolerate acid shock and antibiotic exposure (63). This may explain the distinction between the *rpoB* mutations found in the heat-adapted and control lineages, as heat shock similarly induces a hostile environmental change. The two-component system response regulator *cpxR* was mutated in three lineages in the control arm, but in none of the heat-adapted lineages. Of the three distinct mutations, two were conversions of aspartic acid to asparagine (D190N and D194N) in the OmpR/PhoB-type DNA binding domain (64, 65). This system is involved in controlling the expression of Type II and Type IV secretion system genes and the control of virulence during the switch between exponential and post-exponential phases (66, 67). Prior work in our lab has shown that *cpxR* is necessary for heat shock tolerance (42) – this could underlie an evolutionary inelasticity that favours its mutation in control lineages but not in heat-adapted lineages.

**Table 3:** Selected mutations derived over the course of the ALE determined from Breseq analysis of pooled Illumina short reads. The table includes non-synonymous mutations which emerged in two or more lineages of either the control or heat-adapted protocol branches. All mutations listed were present in at least 25% of pooled sequence reads.

|  | C-1 | C-2 | C-3 | C-4 | C-5 | C-6 | HA-1 | HA-2 | HA-3 | HA-5 | HA-6 |
| --- | --- | --- | --- | --- | --- | --- | --- | --- | --- | --- | --- |
| **pPh38** | Δ37,890<br>bp chr1:<br>173323 | Δ37,890<br>bp chr1:<br>173323 | Δ37,890<br>bp chr1:<br>173323 | Δ37,890<br>bp chr1:<br>173323 | Δ37,890<br>bp chr1:<br>173323 | Δ37,890<br>bp chr1:<br>173323 | Δ37,890<br>bp chr1:<br>173323 | Δ37,890<br>bp chr1:<br>173323 | Δ37,890<br>bp chr1:<br>173323 | Δ37,890<br>bp chr1:<br>173323 | Δ37,890<br>bp chr1:<br>173323 |
| **two-component system**<br>**response regulator LetA** | L68R | L68R | L68R | L68R | L68R | L68R | L68R | L68R | L68R | L68R | L68R |
| **transcription**<br>**termination/antitermination**<br>**protein NusG** |  | R136I |  |  |  | V143D |  |  |  |  |  |
| **DNA-directed RNA**<br>**polymerase subunit beta** |  |  |  | G759E | N1350T |  | G138E |  |  | K924N |  |
| **30S ribosomal protein S10** |  |  |  | Q36K | S102N |  |  |  |  |  |  |
| **hypothetical protein (phasin)** |  |  |  |  |  |  | Δ1bp<br>nt39 | Δ1bp<br>nt39 |  |  | +A 38 |
| **rod shape-determining**<br>**protein MreC** |  |  |  |  |  |  | V266A | V266A |  |  |  |
| **LidA** | Δ1bp<br>nt1566 | Δ1bp<br>nt1122,<br>P330R | E508* | Δ1bp<br>nt1566 | Δ1bp<br>nt1122 | Δ1bp<br>nt1566 |  |  |  |  |  |
| **molecular chaperone HtpG** |  |  |  |  |  |  |  |  | G83E | Q148* | Δ1bp<br>nt503 |
| **rod shape-determining**<br>**protein RodA** |  |  | A128T |  | A128T |  |  | R112S | P332S |  |  |
| **transcriptional regulatory**<br>**protein CpxR** |  | R112L |  | D190N |  | D194N |  |  |  |  |  |
| **ATP-dependent chaperone**<br>**ClpB** |  |  |  |  |  |  |  |  | A478V | A492T |  |
| **ATP-dependent protease**<br>**ATP-binding subunit ClpX** |  |  |  |  |  |  | G202A | E203D |  |  |  |
| **molecular chaperone DnaJ** |  |  |  |  |  |  | F95Y | F95Y | P334S | +C 1100 | R318T |
| molecular chaperone DnaK |  |  |  |  |  |  | M94I | V373L,<br>M94I |  |  | M94I |
| hypothetical protein (mavF) | | | | | | | Δ1bp<br>nt368 | Δ1bp<br>nt368 | Δ1bp<br>nt368 | Δ1bp<br>nt368 | +T 456 |
| NADH-quinone<br>oxidoreductase subunit G |  |  |  |  |  |  |  |  |  | G455D | E768K |

### Central role of the DnaJ/DnaK/ClpB disaggregase complex in heat shock adaptation

Adaptation to heat stress, as expected, took multiple unique but overlapping pathways. Canonical heat shock factors were common targets of mutation, and included *htpG, clpB*, *clpX*, *dnaJ*, and *dnaK*. These are, variously, chaperone and co-chaperone proteins that act on misfolded and aggregated proteins to resolve the effects of protein damage (68). The *bona fide* HSP40 co-chaperone *dnaJ* was mutated in all five heat-adapted lineages, while neither of the HSP40 homologues *cbpA* or *djlA* (69–71) received any mutations. Four distinct mutations were acquired in *dnaJ*, including three missense mutations and a single nucleotide insertion. Three mutations – P334S, R318T, and an insertion at nucleotide 1100 (codon 337) – are in the C-terminal end of the polypeptide. This region has been linked to the binding between DnaJ and DksA, a regulator of RNA polymerase and the stringent response which influences heat shock resistance in *L. pneumophila* (72–75). Autodimerization and chaperone functions in DnaJ also involve this C-terminal region (76). Two lineages derived the same mutation – F95Y – in a linker region connecting the J-domain and central zinc-binding regions of DnaJ. This phenylalanine position is conserved across bacterial and eukaryotic homologues of DnaJ as a residue in the first of three DIF (DVF in *Legionella*) motifs (77). Experimental substitution of this residue in *E. coli* causes a growth defect at low temperatures but has no effect on high-temperature growth or on motility. The integrity of this glycine-phenylalanine rich region is thought to be involved in proper DnaK/DnaJ ATP cycling and substrate release (77). This mutation in DnaJ may contribute to the constitutively high expression of heat shock genes seen in our heat adapted lineages by causing an accumulation of DnaK-substrate complexes and causing a de-repression of steady state *rpoH* mRNA levels (78).

DnaK, a bacterial HSP70 and collaborator with DnaJ in protein refolding, acquired the same mutation in three lineages – M94I. Both F95Y mutations in DnaJ were coincident with this DnaK allele, with HA-2 also possessing a second V373L missense mutation in DnaK. These mutations both alter residues exposed on the AlphaFold predicted surface of the DnaK protein (79, 80), suggesting that their effects lie in modification of the substrate binding or co-chaperone dynamics of the heat shock response network. The DnaK/DnaJ chaperone-co-chaperone system also interacts with ClpB, or HSP100, a AAA+ ATPase with homologue found in all three domains of life (81–83). In various species of bacteria, ClpB assists in the tolerance of high temperatures, ethanol damage, and oxidative stress by acting as a disaggregase of tangled polypeptides (81). Its proper function depends on the assembly of an active hexamer from ClpB monomers, forming a central channel that the misfolded substrate is threaded through. Interaction with DnaK, as well as hexameric assembly of the protein complex, depends on the M-domain of ClpB (84, 85). This section of the larger AAA+ domain includes four surface-exposed α-helices and spans the two mutations observed in ClpB (A478V and A492T). The presumed effect of these mutations is to influence protein-protein interactions either during self-hexamerization or when interfacing with DnaK (86, 87).

### Refinement of the heat shock response by protein degradation

A second hexameric AAA+ ATPase, ClpX, also showed signs of adaptation with mutations in two lineages targeting adjacent residues – G202A and E203D. Unlike ClpB, ClpX favours proteolysis over refoldase activity, acting in concert with the peptidase ClpP (88). The two mutated residues are both in the AAA+ domain of ClpX and may allow for a refinement of the proteolytic capacity of *L. pneumophila* that remediates the enrichment of unfolded and aggregated protein precipitated by high temperature exposure.

As we had previously observed (75), the 90 kilodalton heat shock protein HtpG has an unlikely role in the heat resistance of *L. pneumophila*. Its up-regulation after heat shock, role in high temperature growth in other species, and its chaperone function all strongly support its identity as a heat shock protein (89–91), but the deletion of *htpG* from a wild-type background increases the ability of *L. pneumophila* to survive exposure to 55 °C for 15 minutes. This is further corroborated by the observed mutations in the ALE, with three different mutations observed in three of the heat-adapted lineages. Two of these mutations – Q148* in HA-5 and a single base pair deletion at position 503 in HA-6 – abrogate the structure of the chaperone, leading to the loss of recognition by an anti-HtpG antibody. These three alleles provide the same resistance to heat shock as the full deletion of *htpG*, and do not complement the deletion. We hypothesized that these presumed loss of function mutations may give a growth advantage between heat shock bottlenecks. We tested this in a relative fitness experiment measuring the growth of a deletion mutant heterologously expressing the wild-type and derived alleles of *htpG*. Strains lacking *htpG* were more relatively fit than one expressing wild-type *htpG*. This was also true for all three derived alleles of *htpG*; the loss of HtpG function during the ALE could therefore have promoted both increased growth in AYE and survival under heat shock. As *htpG* mutations were only seen in our heat-adapted lineages, it seems likely that the increase in heat resistance is the more significant factor favouring their selection.

### Mutations in virulence factors

Strikingly, the *lidA* gene was mutated singly or doubly in each of the six control lineages but in none of the heat-adapted lineages. Each lineage acquired either one of two frameshift mutations (single base deletions at nucleotides 1122 or 1566) or a premature stop (E508*). LidA is a Rab GTPase-binding effector protein trafficked into the host cell by Icm/Dot Type IVb secretion system whose role in infection seems largely involve disruption of the host trafficking system (92, 93). The role that these mutations in *lidA* play in this evolutionary system is obscure considering that (i) the ALE was carried out without passage through host cells, (ii) effector proteins are thought to be highly redundant and the repeated mutation of *lidA* is *a priori* unexplained, and (iii) *lidA* mutation seems to have been highly selected for in control lineages but disfavoured during heat-adaptation. The study which characterized *lidA* found that its absence caused the Icm/Dot complex to become toxic to the bacterium, which was attributed to a possible toxic increase of solute permeability (94). An increase of permeability following the loss of LidA may support increased nutrient uptake in AYE, though the factors favouring complete saturation of *lidA* mutations across all the control lineages and none of the heat-adapted ones remain unclear.

Loss-of-function mutations were a recurrent process across both branches of the ALE with lpg0563 (*phaP*) and *mavF* also acquiring frameshift indels in multiple heat-adapted lineages. Each of the five heat-adapted lineages acquired one of two distinct null mutations in *mavF* – a single base pair deletion of nucleotide 368 or a single base pair insertion at position 456. The biological mechanism favouring this mutation is difficult to deduce as the protein is translocated through the Icm/Dot system and therefore likely no *prima facie* relevance to heat shock survival in the absence of host cells (95).

### Future avenues

With hospitals recognizing the dangers that *L. pneumophila* introduces when it contaminates their HWDS plumbing, it is important to understand its life in these systems before it causes nosocomial infection. Heat-based control measures are common, but *L. pneumophila* is often able to survive both routine hot water temperature regimes and remedial superheat-and-flush interventions (23, 96). Our ALE model of *L. pneumophila* microevolution under selective pressure from transient heat shock shows that it can follow several separate evolutionary pathways to acquire heat resistance and replicates *in situ* studies that find increased heat resistance in isolates taken after failed heat treatment (23). *L. pneumophila* lineages in this system evolved in isolation from the broader HWDS microbiome. Even without these exogenous sources of novel DNA, these populations evolved under the pressure of heat shock to survive up to 59 °C. These mutations were concentrated both in well-known chaperones and proteases (*dnaJ*, *dnaK*, *clpX*, etc.), as well as in unexpected proteins with unrelated functions (*phaP*, *mavF*, etc.). This model helps inform our understanding of the forces operating on *L. pneumophila* in hot water plumbing and provides a platform for future work to directly confirm the biological significance of the genetic changes that we have observed in this study.

## Materials and Methods

### Routine culture methods

*L. pneumophila* strains were routinely grown from frozen stocks stored at-80 °C on CYE (ACES-buffered charcoal yeast extract) agar plates (Sigma-Aldrich) adjusted to pH 6.90 with KOH and supplemented with 0.25 g/L L-cysteine and 0.4 g/L ferric pyrophosphate and incubated at 37 °C for 72 hours. Single colonies were sub-cultured into AYE broth (CYE without charcoal or agar) and grown shaking at 37 °C overnight. To simulate residence in tap water, Fraquil, a defined low-nutrient medium (97) that simulates freshwater was used to suspend *L. pneumophila*. *E. coli* strain DH5α was grown in LB agar or broth, as indicated, at 37 °C overnight. Where required, media was supplemented with 5 mg/L (*L. pneumophila*) or 25 mg/L (*E. coli*) chloramphenicol, 25 mg/L kanamycin sulfate and/or 0.1 mM isopropyl β-d-1-thiogalactopyranoside (IPTG). The strains used in this study are listed in Table 1. *Vermamoeba vermiformis* was maintained in modified PYNFH (ATCC media 1034) at room temperature (20-28 °C) and passaged once per week (42).

### Adaptive laboratory evolution procedure

The adaptive laboratory evolution experiment was designed with two arms deriving from one common ancestral strain. *L. pneumophila* str. Philadelphia-1 (ATCC 33152) was obtained from the American Type Culture Collection; it is a clinical isolate collected from the 1976 outbreak of Legionnaires’ Disease in Philadelphia. Because this isolate was sourced from a cooling tower, we expected it to be poorly adapted to growth in laboratory conditions in axenic medium, a known selection pressure in ALE experiments. To account for this, we included both an experimental arm which was exposed to heat stress and a control arm which was not exposed to heat stress (Figure 1A and B) to compensate for these effects.

To begin the experiment, a single culture of Philadelphia-1 was divided into 12 independent lineages to assess the parallelism between adaptive paths, divided into six evolved lineages that were exposed to heat shock and six control lineages that were not. Each population was grown in AYE overnight to stationary phase. Bacteria were triply washed in Fraquil, suspended at OD_600_ 0.1 (approximately 10^8^ CFU/mL) and incubated for 24 hours at room temperature, as in (98). For each suspension, 1 mL was dispensed into a 13 mL polypropylene culture tube (Sarstedt). Tubes of the treated lineage were immersed for 15 minutes in a pre-heated water bath set to the challenge temperature (55°C to 59°C). Samples were withdrawn from the water bath and cooled passively to room temperature. Tubes of the control lineages were left at room temperature. Control lineages were propagated by inoculating 1.5 mL AYE with 15 µL (approximately 1.5 x 10^6^ CFU) of the untreated suspension, to mimic the population bottleneck created by heat shock. Evolved lineages were propagated by inoculating 750 µL of 2x concentrated AYE with 750 µL of the heat-treated suspension. Cultures were grown overnight to stationary phase, as before, and a 100 µL aliquot was stored at-80 °C in AYE + 15% glycerol to maintain a frozen archive of the full population at each passage.

The experiment was continued for 70 passages with the temperature of heat stress increased on an *ad-hoc* basis to maintain the strength of selective pressure. Based on preliminary experiments, the experiment was begun with a 15-minute exposure to 55 °C. After 10 passages under selection, this was increased to 57 °C. This was further increased to 58 °C after 30 passages and finally to 59 °C after 50 passages.

### Heat shock survival measurements

Following the conclusion of the experiment, heat adapted populations were resuscitated from frozen stocks, grown on CYE, and suspended in Fraquil as above. All lineages were tested at five passage intervals for heat shock survival after 20 minutes of exposure to 55 °C, 57 °C, and 59 °C in 60 µL volumes simultaneously in a Veriti 96-well thermocycler and actively cooled to 20 °C, with control samples held at room temperature. CFU counts were determined by serial dilution and plating in 10 µL aliquots on CYE plates.

A similar strategy was used to determine the heat shock tolerance of mutants expressing the various derived alleles of *htpG*. Samples were prepared and aliquoted as above and exposed to 55 °C for 0, 10, 20, and 30 minutes in a Veriti 96-well thermocycler and actively cooled to 20 °C. CFU counts were conducted as above and normalized to the CFU count of the 0 minutes exposure sample.

### Isolate collection and sequencing strategy

Single colonies were isolated from the populations generated after 70 passages and stored at-80 °C. Ten isolates were collected from each heat-adapted lineage and five isolates were collected from each control lineage. In addition, we resequenced our lab stock of Philadelphia-1 to identify any mutations that were fixed in the ancestral population at the outset of the ALE. DNA was collected from these isolates using the Wizard Genomic DNA Purification kit (Promega) and submitted to the McGill Genome Center for library preparation and sequencing using a 600 cycle MiSeq v3 reagent kit.

Fastp (99) with default settings was used to both clean and perform quality control on paired-end reads. Read alignment and variant calling were conducted using Breseq 0.35.4 (100), bowtie2 2.3.5.1 (101, 102), and R 4.0.3 (103) with the NCBI-deposited sequence (CP015927.1) of the ancestral Philadelphia-1 ATCC 33152 strain as reference genome. Because the per-isolate read coverage was low (mean = 14.37, s = 2.37), there were often regions of missing coverage lacking aligned reads to confidently assign allele identity. To increase the depth of coverage, we analyzed pooled per-lineage sequence data by combining the reads together of all ten sampled isolates for each heat-adapted lineage or all five sampled isolates for each control lineage to reconstruct the allele frequencies in the final populations. We re-analyzed these pooled reads using Breseq using the same parameters to generate our final list of called variants.

### qPCR protocol and analysis

RNA was collected from 1 ml samples of *L. pneumophila* suspended at an OD_600_ of 1 either heat shocked at 55 °C for 15 minutes or held at room temperature for the same duration using TRIzol (ThermoFisher Scientific), as previously described. RNA samples were treated with DNase I and Dnase inactivation reagent (ThermoFisher Scientific) before quantification and storage in nuclease-free water (ThermoFisher Scientific) at-20 °C until required. Protoscript II (New England Biolabs) was used to reverse transcribe RNA into cDNA as template for qPCR analysis using the manufacturer’s protocol. No-RT control reactions were produced in the same way with nuclease-free water replacing Protoscript II.

All qPCR experiments were performed in an Applied Biosystems 7500 Fast Real-Time PCR machine using iTaq Universal SYBR-green supermix (BioRad). Efficiencies for qPCR primers (Table 2) were calculated using a serial dilution of gDNA samples, and all were found to be acceptable (90.9-96.8%) (104). 16S rRNA was used as a common housekeeping gene to normalize Ct values (75) and efficiency-adjusted ddCT (105) was used to compare RNA expression levels between bacteria exposed and naive to heat shock.

### Mutant construction

To determine the role of HtpG over the ALE procedure, we constructed merodiploids by supplementing the various derived alleles of *htpG* on a plasmid. Plasmids were constructed on a pXDC39 backbone allowing *htpG* to be expressed under its native promoter. Using Phusion polymerase and primers kpni-htpg-F and xbai-htpg-R, we amplified the treated lineages T3 (*htpG*^G83E^), T5 (*htpG*^Q148*^), and T6 (*htpG*^Δ1bp 503nt^) alleles from the purified gDNA of single isolates from passage 70. These fragments, carrying the gene sequence and its upstream region, were ligated into pXDC39 to produce plasmids p*htpG*^G83E^, p*htpG*^Q148*^, and p*htpG*^Δ1bp 503nt^. These plasmids were electroporated into a previously constructed KS79-based *htpG* deletion mutant (75) to assay for relative fitness and to isolate the derived *htpG* alleles onto a wild-type background.

### Direct fitness competition in AYE

To determine the relative competitive fitness of *L. pneumophila* before and after our ALE model, we constructed strains that could be co-cultured in AYE and distinguished by the presence or absence of GFP. The GFP-negative control vector pMMB207c and the GFP-positive pXDC31 are distinguished by the insertion of an GFP coding sequence whose expression is driven by pTac and ultimately induced by IPTG. Both plasmids were introduced into the wild-type ancestor Philadelphia-1 and the derived lineages C-1 and HA-1 by electroporation, as previously described (75). A single isolate was collected from each electroporation to represent the fitness of the population in competition.

Strain pairs containing these two plasmids were separately grown overnight and normalized to an OD_600_ of 0.1 in AYE supplemented with chloramphenicol for use in a direct competition assay, described in brief below. Competitions were carried out under standard incubation conditions. For each replicate, 500 µL of each culture of the strain pair were combined at a 1:1 ratio. A 100 µL sample of this mixture was used to inoculate 900 µL AYE with chloramphenicol and adjusted to a final OD_600_ of 0.01 and incubated overnight. The overnight culture was diluted to an OD600 of 0.01 and used to continue the assay in the same way for a total of three transfers. CFU concentration for each timepoint was counted by serial dilution plating on AYE with chloramphenicol and IPTG. Population fractions were manually counted under a UV black light to distinguish between fluorescent and non-fluorescent colonies.

### Host cell infection

*Vermamoeba vermiformis* was used as a permissive host to assess the infectivity of *L. pneumophila*. The amoeba was routinely grown in 25 mL vented flasks (Sarstedt, Corning) at room temperature and passaged twice a week in modified PYNFH media with FBS and buffer. Amoeba populations were expanded in 75 mL flasks 3 days before infection and harvested by centrifugation at 200 × *g* for 10 minutes.

Host cells were normalized in modified PYNFH without FBS or buffer at 5 x 10^5^ cells/ml and 1 mL was added to each well of a 24-well cell culture plate. Infectious bacteria were collected from fresh colonies on CYE plates and normalized in Fraquil to an OD of OD_600_ 0.01 (approximately 10^7^ CFU/mL). To start infections with an MOI of 0.1, 5 µL bacterial suspension was added to each well and the initial CFU count was conducted on the gently mixed supernatant. Growth was carried out for 3 days at 37 °C with CFU counts performed daily.

### Western blotting

Protein samples for Western blotting were prepared using a modified version of the Bio-Rad electrophoresis guide (42). Briefly, *L. pneumophila* pellets were harvested by centrifugation from a 200 µl sample of overnight culture rinsed in Fraquil and normalized to an OD_600_ of 1. Pellets were homogenized by bath sonication (Cole-Parmer) on ice for 10 minutes after thorough suspension in 200 µl sample solubilization buffer. Samples were diluted with 250 µl 2X Laemmli sample buffer (BioRad) and held at room temperature for 20 minutes. The sample was centrifuged for 30 minutes, and the protein sample was collected from the resulting supernatant.

The proteins were migrated in hand-cast 12.5% polyacrylamide gels (BioRad) at 50 V for 30 minutes and 120 V for 90 minutes. Separated proteins were transferred onto polyvinylidene difluoride (OVDF) membranes (BioRad) at 25 V for 12 hours at 4 °C in preparation for immunoblotting and blocked using TBS wash buffer with 5% skim milk powder. Ab225994 (rabbit anti-*E. coli* HtpG; AbCam) was found to be cross-reactive to *L. pneumophila* HtpG and was used as the primary antibody at a 1:5000 dilution, along with rabbit anti-IcdH (Sigma-Aldrich) (1:10000) as a loading control. Primary incubation was carried out for 2 hours, followed by rinsing and blotting with a 1:5000 goat anti-rabbit HRP (Sigma-Aldrich) for 1 hour. Bands were developed using a ECL Prime Western Blotting Detection Reagent (Cytiva BioSciences) and imaged using a ChemiDoc MP Imaging System.

## Data Availability

Raw read data from sequenced isolates collected after 30 and 70 passages have been deposited to the National Center for Biotechnology Availability’s sequence read archive and are available under BioProject Accession number PRJNA956983.

## Acknowledgements

Many thanks to Jesse Shapiro for providing guidance on ALE. This work was supported by a Natural Science and Engineering Research Council of Canada Discovery Grant (RGPIN/04499-2018) to SPF. JL was the recipient of a Canada Graduate Scholarship—Master’s award from the Natural Science and Engineering Research Council of Canada.

## Notes

### Competing Interest Statement

The authors have declared no competing interest.

### Summary of Updates

Include supplementary materials that was not transferred by AEM.

